# Molecular dynamics descriptors for 1,079 post-translationally modified protein systems

**DOI:** 10.64898/2026.08.23.746532

**Authors:** Kaining Liu, Qiuting Qian, Jiahua Peng, Dongge Ma, Yifei Yao, Jingjing Zhao, Ying Chi

**Affiliations:** Department of Pharmacy of the Second Affiliated Hospital of Zhejiang University School of Medicine, and Zhejiang University–University of Edinburgh Institute (ZJE), Zhejiang University, No. 866 Yu Hang Tang Road, 310058, Zhejiang Province, China; Edinburgh Medical School: Biomedical Sciences, College of Medicine and Veterinary Medicine, The University of Edinburgh, United Kingdom

## Abstract

Databases of post-translational modifications (PTMs) catalogue modified sites and increasingly add static structural context, but trajectory-derived descriptors remain scattered across specialised tools and general molecular dynamics archives. Dyna-MO PTM brings together 1,079 AlphaFold 3-seeded systems covering lysine acetylation, lysine and arginine monomethylation, and serine, threonine and tyrosine phosphorylation. Each system is linked to three completed 10 ns replicas generated with CHARMM36m and TIP3P, for 32.37 μs of aggregate sampling. A 118-column table joins simulation and quality-control provenance with global relaxation measures, site solvent exposure, rotamers, secondary structure and ionic-contact proxies. Versioned identifiers connect the records to starting structures, trajectories, manifests and analysis scripts. Researchers can use the resource to filter PTM contexts, reproduce descriptors, prioritise longer simulations and evaluate trajectory-analysis or generative methods. The trajectories describe finite-window relaxation rather than equilibrium free energies, kinetics or matched PTM effects.

## Background & Summary

Post-translational modifications create distinct protein states by changing local charge, hydrogen-bonding capacity or steric geometry. Large annotation resources catalogue these sites across the proteome^1^. Linking site annotations to structural and trajectory-derived descriptors can guide system selection and comparative analysis. Short simulations complement static annotations with local relaxation information.

PhosphoSitePlus, dbPTM, PTMcode, ActiveDriverDB and UniProt provide complementary site-level curation^2–6^. Structural resources supply a different view. The Protein Data Bank defines modified chemical components, whereas AlphaFold 3 accepts PTM-bearing inputs^7,8^. StrucPTM reports experimentally observed structural contexts, and curated rotamer libraries describe static PTM side-chain preferences^9,10^. General MD resources, including ATLAS, mdCATH, GPCRmd and MDRepo, provide standardised, specialist or community-deposited trajectories^11–14^. PTMdyna and Vienna-PTM instead provide PTM-aware analysis or system-preparation workflows^15,16^. Among these representative resources, none is organised around the same combination of multi-class PTM chemistry, protein-level trajectories, cross-system descriptors and file-level provenance (Table 4).

Dyna-MO PTM provides that combination for acetyl-K, monomethyl-K, monomethyl-R, phospho-S, phospho-T and phospho-Y. It maps two production routes into a shared 118-column schema, with three completed 10 ns replicas per released system. Simulations use the CHARMM36m parameter family with TIP3P water^17,18^. The 738 K/R systems were generated for this resource. The other 341 systems come from the DynaMo-phos phosphorylation archive described with ProteinFlux^19^. Their provenance, quality-control rules and replica aggregation remain route-specific within the common schema. The primary contribution is the alignment of six PTM classes, protein trajectories, descriptors and traceable files under versioned identifiers. Figure 1 defines the released collection, and Figure 2 traces its construction and data products.

**Figure 1.**
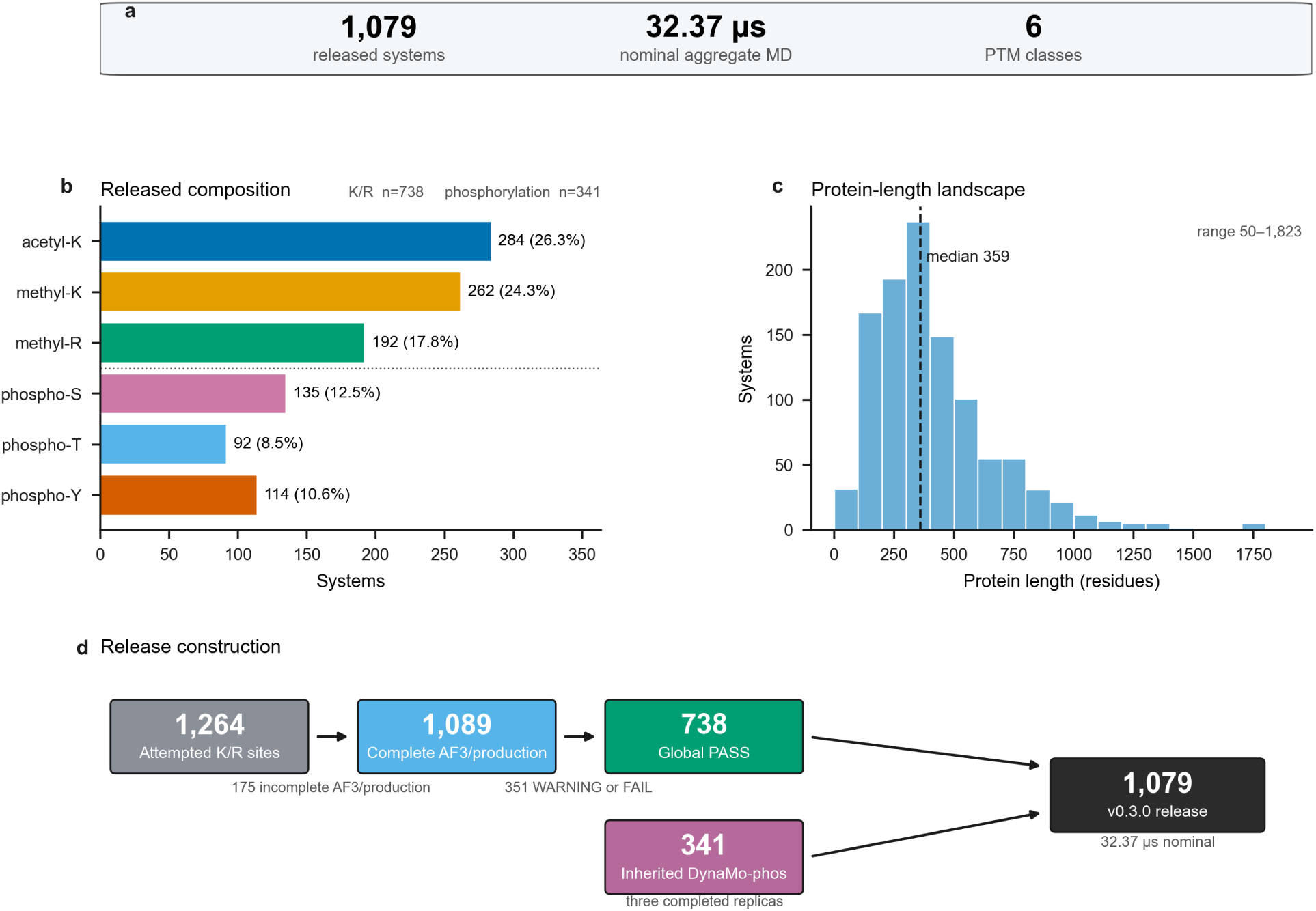
Dyna-MO PTM v0.3.0 dataset overview. (a) The dataset comprises 1,079 systems, 32.37 μs of completed MD sampling and six PTM classes. (b) Counts by modification class separate the 738-system K/R subset from the 341-system DynaMo-phos subset. (c) The protein-length distribution has a median of 359 residues and a range of 50–1,823. (d) The K/ R retention funnel contains 738 systems from 1,264 attempted sites. The 341 phosphorylation systems were inherited under the DynaMo-phos completion rule and did not pass through the K/R funnel; together the two strata form the 1,079-system release.

**Figure 2.**
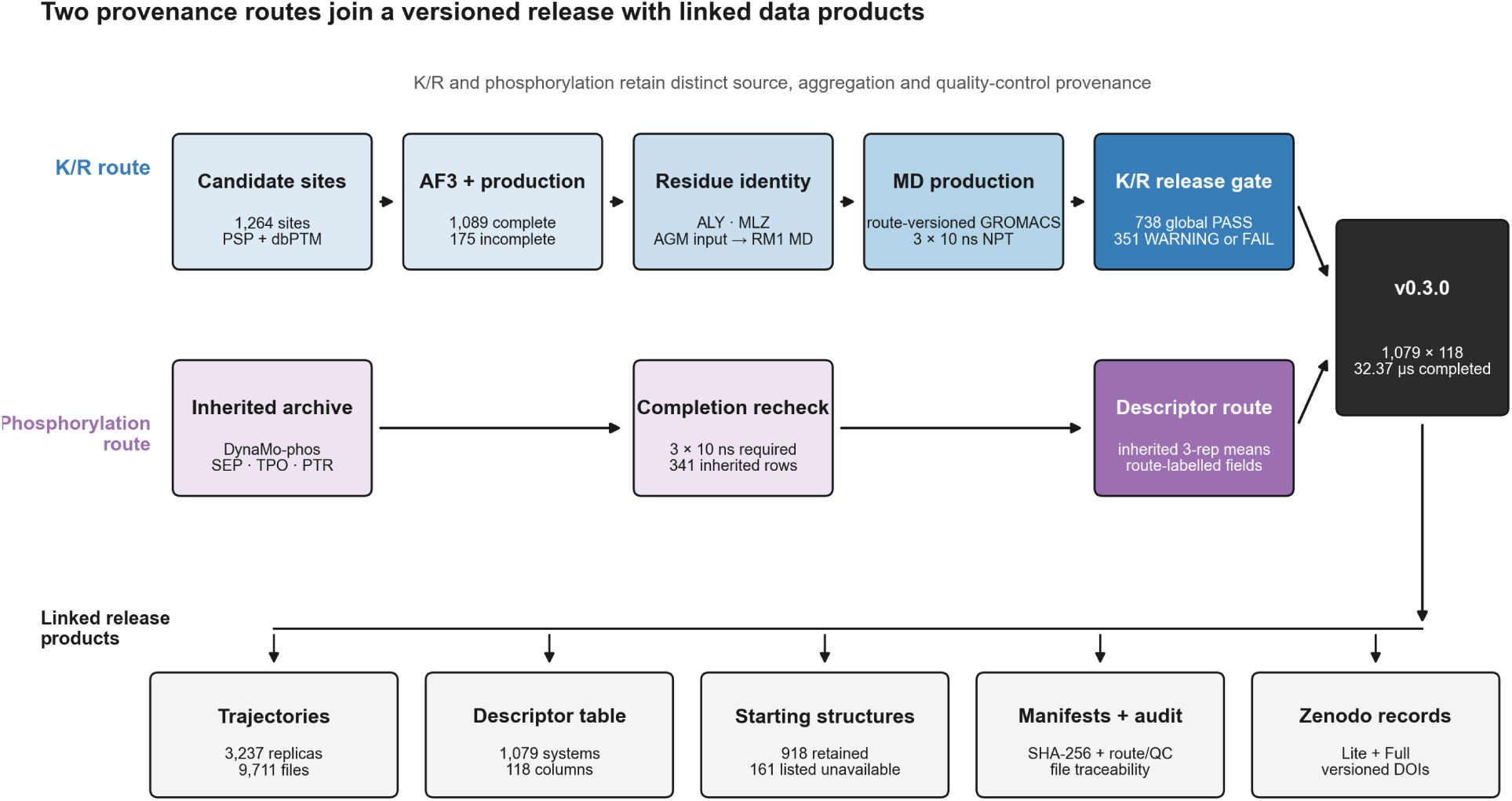
Construction pipeline and released data products. The K/R route links curated candidate sites to AF3 inputs, route-versioned MD production and the global release gate. ALY and MLZ retain the same input and MD component names. AGM identifies the rMMA input component, whereas MD topology generation uses RM1 after deterministic connectivity conversion. The phosphorylation route contributes 341 inherited DynaMo-phos rows and route-labelled three-replica descriptors. Its observed-file recheck confirms three completed 10 ns replicas for every released system. Both routes join the 1,079 × 118 v0.3.0 schema and its linked products: 3,237 trajectories, 918 retained starting structures, file manifests and audits, and the Lite and Full Zenodo records. The 161 unavailable batch-2 phosphorylation inputs are listed explicitly.

The release is intended for descriptor filtering, reproducible trajectory reanalysis, method development and selection of systems for longer simulations. Because most records lack matched unmodified controls, differences among PTM classes characterize the selected resource rather than effects of modification.

## Methods

### System curation and starting structures

Initial modified-residue sites came from local PhosphoSitePlus and dbPTM aggregations. The surviving records lack database snapshot dates, evidence filters and conflict-resolution rules, so the upstream sampling frame cannot be reconstructed exactly. The released rows form a curated convenience sample rather than a representative census of proteome-wide PTM sites.

Every released row retains a versioned system identifier, UniProt accession, modified residue and site number. The 1,264-row K/R ledger also records candidate identifiers, truncation coordinates, selected AF3 samples and source PDB paths. Phosphorylation rows retain their DynaMo-phos batch and UniProt-site mapping. Together, these fields identify the final sampling units.

The input set comprised 1,264 K/R sites represented in AlphaFold 3 by ALY, MLZ or AGM, and 343 phosphorylation sites represented by SEP, TPO or PTR. These names identify components in the input structures; the MD residue definitions are recorded separately. K/R starting structures were generated as single-chain predictions with a locally modified derivative of AlphaFold 3 v3.0.0 (official tag commit `4f52a3bb624fd7245fb1ad0bf62d752ffa46f0ce`). Each input JSON used chain A, model seed 1 and a CCD modification at the target sequence position. The data pipeline and inference stages were both enabled. Protein searches used small BFD, MGnify 2022_05, UniProt 2021_04 and UniRef90 2022_05; template searches used the local PDB mmCIF collection and PDB seqres 2022_09_28. Inference otherwise used the packaged v3.0.0 model configuration, with the Triton attention implementation, and generated five diffusion samples per input. The retained sample maximised mean pLDDT within the selected structural core rather than the AF3 ranking score. This rule aligned sample selection with the region retained for simulation; it was not intended to re-rank the complete AF3 output. The archived runner, input generator and sample-selection scripts have SHA-256 values `28b00cc1…60a24`, `cc2cdef0…5087` and `01f240bc…af0d`, respectively.

Before MD, the pipeline retained the largest contiguous window that contained the site and had per-residue pLDDT of at least 70. A system was excluded if this window contained fewer than 50 residues or had mean pLDDT below 70. Truncation exclusions and production failures account for the 175 incomplete K/R systems in Figure 1d. The phosphorylation subset came from DynaMo-phos batches 2 and 3. Per-residue pLDDT was stored in the PDB B-factor column. The reconciled archive contains 918 modified starting PDBs; Data Records lists the 161 unavailable batch-2 inputs.

### PTM installation, technical correction and force-field provenance

Topologies were built with the July 2022 GROMACS CHARMM36 force-field port distributed by the MacKerell Lab. Distributed residue definitions were used for ALY, MLZ, SEP, TPO and PTR. The source AGM geometry placed its methyl substituent on Cδ and therefore represented 5-methylarginine rather than biological *N*^G^-monomethyl-L-arginine (rMMA). Before rMMA topology generation, the 208 source structures were deterministically converted to residue RM1, with the methyl carbon bonded to a terminal guanidino nitrogen. The RM1 definition combines the distributed CHARMM36m 2MR residue scaffold with MGUA guanidinium-head atom types and partial charges. No new atom type or bonded or non-bonded parameter was fitted. The conversion, residue charge (+1), connectivity and minimum methyl-environment distance are checked before production. The parameter lineage follows prior CHARMM modified-residue work^20^. The exact distributed files, checksums and project-authored conversion script are archived in the lite record described below.

Hydrogens were rebuilt with `gmx pdb2gmx -ignh`. Standard charged termini were selected, NH_3_^+^ at the N terminus and COO^−^ at the C terminus; no terminal caps were added. This treatment also applies to boundaries introduced by structural-core truncation. Histidine protonation followed the CHARMM36m `pdb2gmx` automatic assignment because the preparation command did not request interactive histidine selection and no per-system override was supplied.

The checks establish topology completeness and internal consistency for the recorded production route. RM1 was not subjected to a QM refit or experimental conformational and thermodynamic validation, so applications beyond this protocol require a separate force-field assessment.

Each protein was placed in a rhombic dodecahedron and solvated with TIP3P water. The preparation script first used 0.90 nm protein-to-box padding. If the solvated system exceeded the operational ceiling of 500,000 atoms, it was rebuilt at 0.75 nm and, when still necessary, at 0.60 nm. Among the 738 released K/ R systems, 731 used 0.90 nm, five used 0.75 nm and two used 0.60 nm. Sodium or chloride counterions neutralised the net charge. The `genion` command used `-neutral` without `-conc`; no bulk salt concentration was imposed. The selected padding and ion counts are retained in per-system preparation logs.

### Energy minimisation, equilibration and production

Energy minimisation used steepest descent for at most 50,000 steps, with a 1,000 kJ mol⁻¹ nm⁻¹ tolerance and 0.01 nm initial step. Equilibration began with 100 ps NVT at 310 K using the velocity-rescaling thermostat^21^ (τ_t_ = 0.1 ps). It continued with 200 ps NPT at 310 K and 1.0 bar using the Parrinello–Rahman barostat^22^ (τ_p_ = 2.0 ps). Production targeted three 10 ns replicas. It used the NPT ensemble, velocity rescaling (τ_t_ = 0.5 ps) and Parrinello–Rahman pressure coupling (τ_p_ = 5.0 ps). The remaining settings were a 2 fs integration step, LINCS constraints on hydrogen-containing bonds^23^ and the Verlet cutoff scheme. Smooth PME electrostatics^24^ used 1.2 nm Coulomb and Lennard-Jones cutoffs. Coordinates were saved every 100 ps, producing 101 frames for a completed 10 ns trajectory.

The legacy production templates requested fresh velocities with `gen_vel = yes` and `gen_seed = -1`. The regenerated RM1 route instead records deterministic, distinct `gen_seed` and velocity-rescaling `ld_seed` values for every system and replica. Legacy production records identify GROMACS 2026.0, whereas the audited RM1 regeneration uses GROMACS 2025.2^25^. Software version and seed fields are retained separately for each production route.

### Quality control gating

Trajectories were post-processed (`gmx trjconv -pbc mol -center`) to remove periodic boundary jumps and centre the protein. The exact legacy global release gate for the K/R subset used GROMACS rep-1 series as implemented in `scripts/md_analysis/traj_qc_ptm.py`. The RMSD inputs were the mean and population standard deviation (`ddof = 0`) over the final 20% of frames and the absolute linear slope over the full trajectory. The Rg input was the full-trajectory coefficient of variation, also calculated with the population standard deviation. Per-residue Cα-RMSF (`gmx rmsf -res -fit`) was a released descriptor rather than a scored gate quantity. Only the number of RMSF residue rows selected the length-dependent RMSD threshold band.

The primary K/R columns follow this GROMACS rep-1 route. `rmsd_equil_mean_A` and `rmsd_equil_std_A` use frames 80 to 100 inclusive, the final 21 of 101 frames, with `ddof = 0` for the standard deviation. `rmsd_slope_A_per_ns`, `rg_cv_percent` and `rg_slope_A_per_ns` use all 101 frames. The site RMSD mean uses the same final-20% window, whereas its slope uses the full series. Separate three-replica fields were calculated from `traj_stats_per_rep.csv`. `rmsd_equil_mean_3rep_A` averages final-50% means, while `rmsd_mean_3rep_A` and `rg_mean_3rep_A` average full-trajectory means. These three-replica summaries serve a different purpose from the rep-1 release-gate quantities.

The retained acetyl-K and methyl-K rep-1 source trajectories were later appended from 10 ns to 100 ns. All release-gate and descriptor values in the master table were generated from the original 101-frame, 0– 10 ns analysis files before that extension. The release payload comprises validated 0–10 ns derivatives linked to the preserved 10 ns TPR and `md_10ns.gro` files, while the appended trajectories remain separate extension data.

The thresholds in Table 1 define the exact legacy global gate. They are pipeline-engineering heuristics inherited from the original K/R workflow, not universal criteria for structural stability. A system is FAIL if any failure threshold is exceeded, WARNING if no failure threshold is exceeded but at least one warning threshold is exceeded, and PASS otherwise. We therefore repeated the gate after multiplying all absolute RMSD, slope and Rg-CV warning and failure cutoffs by 0.8 or 1.2 while holding the dimensionless RMSD relative-fluctuation cutoff at 0.30. The 94 completed methyl-R trajectories that were not regenerated with corrected RM1 chemistry remained ineligible at every scale. Among the 995 chemistry-eligible systems, the number classified PASS was 670 at 0.8×, 738 at 1.0× and 799 at 1.2×. The complete system-level audit is supplied with the analysis scripts.

**Table 1.**
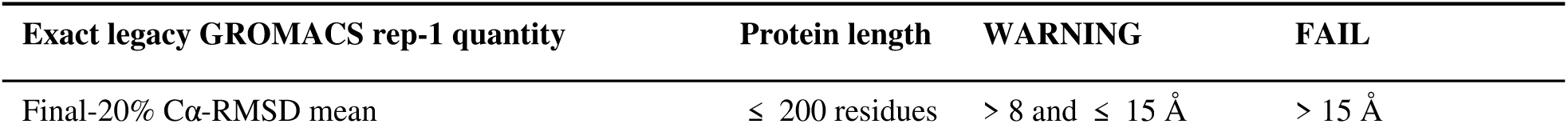
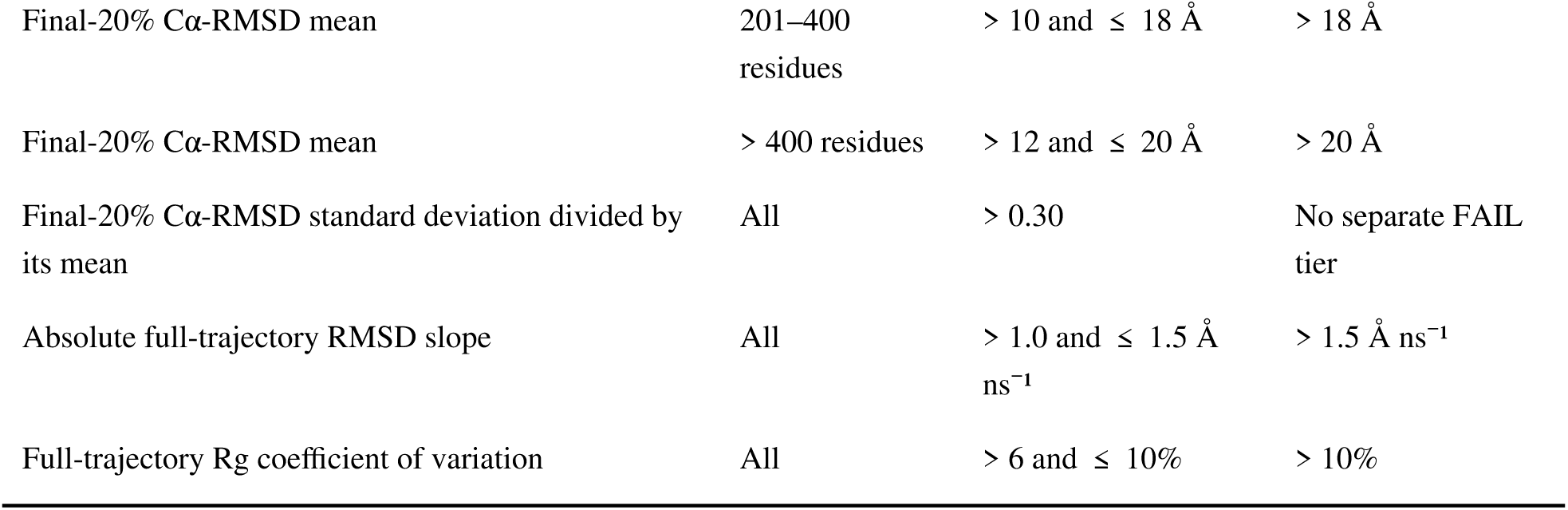
Exact legacy GROMACS rep-1 release-gate thresholds for K/R systems.

The site label is a separate diagnostic based on the final-20% mean Cα-RMSD of a self-aligned ±5-residue window. Values ≤ 3 Å are `SITE_OK`, values > 3 and ≤ 6 Å are `SITE_WARN`, and values > 6 Å are `SITE_BAD`. A `SITE_WARN` label does not exclude an otherwise globally passing system. When global `status` is `PASS`, `final_flag` is `PASS` for `SITE_OK` or `SITE_WARN` and `PASS_SITE_BAD` for `SITE_BAD`; otherwise `final_flag` retains the global `WARNING` or `FAIL`. The release source is the global-PASS set. A stricter companion set additionally excludes `SITE_BAD`. Both sets contain the same 738 K/R systems in v0.3.0 because no globally passing system is `SITE_BAD`. The local self-alignment reduces coupling to global protein displacement.

Of the 1,264 prepared K/R-PTM systems, 738 satisfy the release criteria in v0.3.0 (284 acetyl-K, 262 methyl-K and 192 methyl-R). The corrected RM1 quality-control reconstruction classifies the other 16 completed methyl-R systems as `WARNING`; none is `FAIL`. The phosphorylation subset was inherited under a source-route rule requiring three completed 10 ns replicas. The single-residue placeholder `c9j7s5_y1_phospho` and `o75832_s25_phospho`, whose rep-2 production trajectory was shorter than 1 ns, were excluded. A pre-deposition audit also found that `o75792_s18_phospho` rep 2 ended at 3.8 ns because the inherited parser excluded only trajectories shorter than 2 ns. We resumed the replica from its preserved checkpoint in an isolated copy and completed the planned 10 ns trajectory. The 101-frame analysis trajectory reproduced the inherited processing route, and all dependent three-replica descriptors were recomputed. A second hash-chain ledger records the 23 corrected row fields while preserving the original trajectory and pre-repair master as evidence. The released phosphorylation set therefore remains 341 systems, with the two original exclusions recorded in `phosphorylation_exclusions_v0.3.0.csv`.

For the cross-route validation companion, we recalculated backbone-fitted Cα-RMSD, the full-series RMSD slope and protein Rg from every deposited replica with one GROMACS workflow. File hashes and the 101-frame 0–10 ns time axis were verified before the exact K/R thresholds were replayed on all 3,237 trajectories. The resulting labels support replica-level filtering; they do not replace the historical release rules.

### Per-system descriptor extraction

Analysis windows are field-specific. The K/R rep-1 global and site-RMSD windows follow the release route defined above, and `site_rmsf_mean_A` is the rep-1 mean across a ±3-residue window. K/R rep-2 site-biophysics descriptors and the equilibrium RMSD component of the `_3rep` block use the final 50% of frames. The three-replica RMSD and Rg means use their full series. The phosphorylation primary global fields and populated site-biophysics descriptors were inherited as three-replica means. Rep-1 site-RMSD, site-RMSF and canonical per-class DSSP percentage fields are not available for those 341 rows and remain blank.

Site SASA follows the source-specific analysis route and represents the sum over all atoms of the modified residue. Legacy acetyl-K and methyl-K values used `gmx sasa` on rep 2, with the Protein surface group, a modified-residue output selection and the final 50% of frames. Corrected methyl-R values use MDTraj Shrake-Rupley SASA on a macromolecule-only rep-2 trajectory over the same window. Phosphorylation SASA values are inherited three-replica means from the source route. All values are reported in nm²^26^. The K/R-D/E ionic-contact proxy counts cationic-nitrogen to anionic-oxygen pairs within 4.0 Å across the protein. Legacy acetyl-K and methyl-K values used `gmx mindist` on the final 50% of rep 2. The corrected RM1 route used MDTraj on the same replica and window and included LYS, ARG and RM1 cationic nitrogens and ASP/GLU anionic oxygens. Phosphorylation values are inherited three-replica means. Cross-route interpretation should therefore retain the recorded source implementation.

Residue-specific dihedrals provide χ₁ for all six modification classes and χ₂ for acetyl-K, methyl-K, methyl-R and phospho-Y. The source routes differ only at exact boundary values. Legacy labels use g+ for 0° ≤ χ < 120°, t for χ ≥ 120° or χ < −120°, and g− for −120° ≤ χ < 0°. Corrected RM1 labels use g+ for 0° < χ ≤ 120°, t for χ > 120° or χ < −120°, and g− for −120° ≤ χ ≤ 0°. The distinction affects only an angle reported as exactly 0° or +120°; route provenance is retained for that reason. SASA, contacts, rotamer occupancies and dominant secondary-structure labels are empirical summaries of the recorded finite windows rather than equilibrium-state populations.

The cross-route comparability contract is summarised in Table 2 and expanded in the column dictionary.

**Table 2.** Cross-route comparability contract for trajectory-derived descriptor families.

| Descriptor family | K/R aggregation | Phosphorylation aggregation | Cross-route use |
| --- | --- | --- | --- |
| rmsd_mean_3rep_A,<br>rmsd_equil_mean_3rep_A,<br>rg_mean_3rep_A | Three-replica aggregate | Alias of inherited three-replica aggregate | Preferred common summary, with source implementation still recorded |
| Primary global RMSD, Rg and release flags | GROMACS rep 1 | Inherited three-replica summary or completion field | Route-specific only |
| RMSF fields | Rep-1 and K/R-only three-replica variants | Inherited three-replica primary value | Not harmonized across all six groups |
| SASA, contact, rotamer and DSSP fields | Rep 2, with GROMACS or MDTraj by K/R subtype | Inherited three-replica summary | Route-specific and descriptive |

Modification-site secondary structure was calculated with `gmx dssp`^27^. Codes were collapsed to H (H, G or I), E (E or B) and C (turn, bend or coil). For populated K/R rows, `site_ss_stability` is the percentage of analysed frames assigned to the dominant class. K/R structural-context fields were calculated from the AF3 input. They include the distance to each terminus and the Cα-neighbour count within 8.0 Å after excluding five adjacent sequence positions. The fields also include the mean B-factor of the neighbouring Cα atoms.

### AF3 raw-output confidence extraction

Raw AF3 outputs (`summary_confidences.json` and `confidences.json`) were recovered from the original cloud archives. They were matched exactly to the retained starting PDBs for all 738 released K/R-PTM systems. For 75 batch-3 phosphorylation systems, the best sample was selected by maximum `ranking_score` across five seeds.

A further 105 phosphorylation batch-3 systems were recovered after correcting for a pre-MD truncation offset. The master table uses the post-truncation site index, whereas the archived predictions use original UniProt numbering. Each offset was checked against `truncation_plan.csv`, and positional PAE metrics were calculated at the corresponding original site. `af3_source` records whether the match was direct or offset-corrected. Raw outputs were unavailable for the remaining 161 phosphorylation batch-2 rows, so `af3_*` fields are null for those systems.

### Within-resource paired baseline

For the reverted-to-canonical diagnostic, six phosphorylation systems were selected with two systems from each residue class. The set was fixed before descriptor calculation. In each starting structure, the four phosphate atoms (P, O1P, O2P and O3P) were removed. HETATM records were converted to ATOM records, and SEP, TPO or PTR was renamed SER, THR or TYR. Hydrogens were rebuilt with `gmx pdb2gmx -ignh`, and the resulting unmodified systems were processed with the same minimisation, equilibration and three-replica production workflow. The resulting six-pair set provides a within-pipeline diagnostic; its size and convenience sampling limit inference about phosphorylation effects.

### Extended-trajectory diagnostic

Rep 1 of each of the 284 retained acetyl-K systems was extended to 100 ns with the same production settings. The diagnostic compares the non-overlapping 0–10 ns and 90–100 ns blocks, which have equal duration. For backbone RMSD and Rg, the analysis reports the early and late means, paired difference and RMSE. It also reports Pearson and Spearman correlations and a percentile-bootstrap 95% confidence interval from 20,000 resamples. RMSD is measured relative to frame 0. Because no equivalence margin was prespecified, the analysis quantifies early-to-late agreement without testing equivalence or convergence.

### Statistical summaries used for validation

The Data Overview reports descriptive per-class medians, interquartile ranges and percentile-bootstrap 95% confidence intervals from 10,000 resamples.

Aggregation sensitivity is assessed within the 738 K/R systems by comparing primary rep-1 values with explicit three-replica summaries. Pearson and Spearman correlations and absolute paired differences quantify agreement. The reverted-to-canonical diagnostic uses paired differences and the Wilcoxon signed-rank test after three-replica averaging. With six pairs, the Wilcoxon results are reported as sensitivity estimates rather than evidence of equivalence. The extended-trajectory diagnostic uses paired differences, RMSE, Pearson and Spearman correlations, and percentile-bootstrap 95% confidence intervals from 20,000 resamples.

### Use of generative AI tools

OpenAI Codex assisted with code review, reproducibility audits and language editing. The authors executed and inspected the resulting scripts and statistical outputs against the release artefacts and remain responsible for the text, code and scientific claims.

## Data Records

The v0.3.0 dataset comprises three linked records, joined by the same system identifiers. Version v0.3.1-lite denotes the current Zenodo archive package; the descriptor schema, master table and trajectory payload remain v0.3.0. A starting structure is the modified coordinate model used to initiate MD. A trajectory is one deposited 0–10 ns coordinate series. Per-replica diagnostics describe an individual trajectory under the uniform observed-file checks, whereas per-system descriptors summarize the replicas according to the recorded production route. Provenance records connect each system to its source, processing route, availability status and file checksums.

### Descriptor table and analysis scripts

The source repository contains the descriptor table and analysis scripts^28^. The same release code is archived in the Zenodo lite record. Its MIT license covers repository code, not the deposited data or third-party inputs. The principal table, `submission/master_table_all_v2.csv`, contains 1,079 rows and 118 columns under schema v0.3.0. Companion files provide column definitions, integrity reports, comparability notes, input and exclusion ledgers, the upstream-provenance audit, starting-structure manifests, analysis scripts and figures. The field-level companion `descriptor_compatibility_v0.3.0.csv` maps every master-table column to its descriptor family, value role, availability, aggregation source, pooling policy and missingness policy. The separate `af3_descriptor_associations_v0.3.0.csv` reports the prespecified starting-confidence associations described in Technical Validation. The validation bundle additionally contains 3,237-row replica and 1,079-row system diagnostic tables, the K/R threshold-sensitivity ledger, the reduced-padding periodic-image audit and artificial-terminus distances.

### Trajectory data

The trajectory payload is available in the companion Zenodo record at https://doi.org/10.5281/zenodo.22049767^29^. It covers 1,079 systems and 3,237 completed 10 ns replicas. Each replica contributes a compressed GROMACS `.xtc` trajectory, a `.tpr` topology companion and a `.gro` reference frame, for 9,711 files in total. The payload occupies 133,700,022,208 bytes (133.70 GB), and every trajectory contains 101 frames sampled from 0 to 10 ns at 100 ps intervals. `full_trajectory_manifest_v0.3.0.csv` records the relative path, byte count, SHA-256 checksum, atom count, time range, processing route and payload partition for each file. The K/R partition contains full-system trajectories. The phosphorylation partition contains fitted protein-plus-SEP/TPO/PTR trajectories with atom-matched topologies and reference coordinates. These subset TPR files support topology-aware analysis but do not contain restart state. A second hash chain records the repair and descriptor recomputation for `o75792_s18_phospho`. Payload construction and the independent file audit passed, supporting the measured total of 32.37 μs.

### Retained starting structures

The file `dyna-mo-ptm-v0.3.0-starting-structures.tar.gz` packages 918 retained PDBs. Each structure is matched to a release identifier and checked for the expected input component at the modified site. Coverage includes all 738 K/R systems and 180 phosphorylation systems from batch 3. The 161 unavailable batch-2 starting PDBs are listed in `starting_structure_missing_v0.3.0.csv`. Source paths, SHA-256 checksums and site-validation counts appear in `starting_structure_manifest_v0.3.0.csv`. A separate quarantine ledger records the 16 corrected methyl-R systems excluded by the reconstructed global gate. The companion Zenodo lite record at https://doi.org/10.5281/zenodo.22059934^30^ combines the table, retained structures, manifests, figures and analysis scripts. Raw trajectories belong to the full record.

### Web interface

The v0.3.0 web build has been generated from the repaired master table and deployed at https://kkkniengliu.github.io/dyna-mo-ptm-web/. An unauthenticated synchronization audit confirmed 1,079 system summaries, 48 fields per summary and 918 retained starting structures against the locked local release. Figure 4 documents this deployed build. The static API follows schema v0.3.0, which is declared by `/api/descriptor-compatibility.json`; it also exposes `data/master_table_all_v2.csv`, an index at `/api/systems.json` and per-system records at `/api/system/<system_id>.json`.

**Figure 3.**
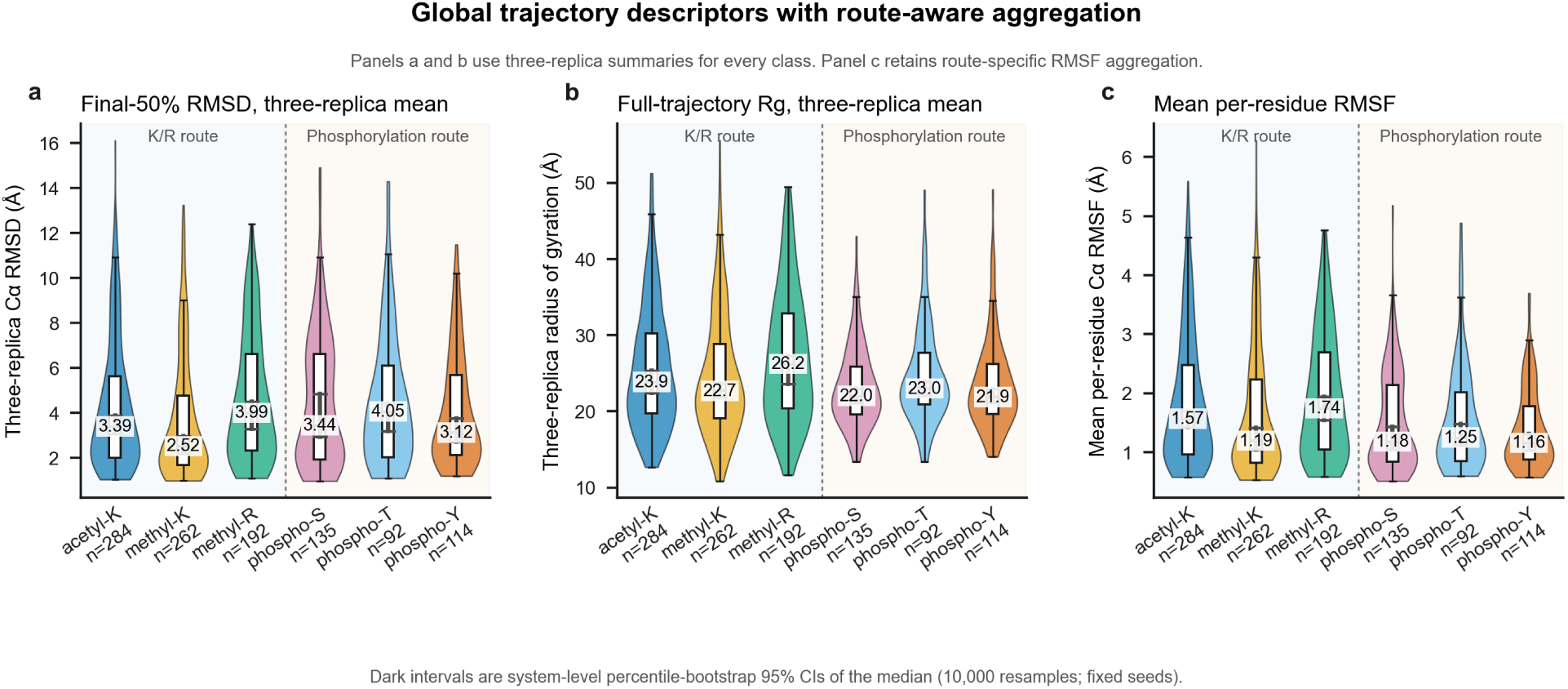
Global trajectory descriptors across six PTM types with the two production routes shown as background strata. Violin plots show (a) the three-replica mean of final-50% backbone-aligned Cα-RMSD, (b) the three-replica mean of full-trajectory Rg and (c) mean per-residue Cα-RMSF. Dark intervals are system-level percentile-bootstrap 95% CIs of the median from 10,000 resamples with fixed seeds. Panels a and b use three-replica summaries for every class. Panel c retains route-specific aggregation because no harmonised three-replica RMSF field is available across all six groups.

**Figure 4.**
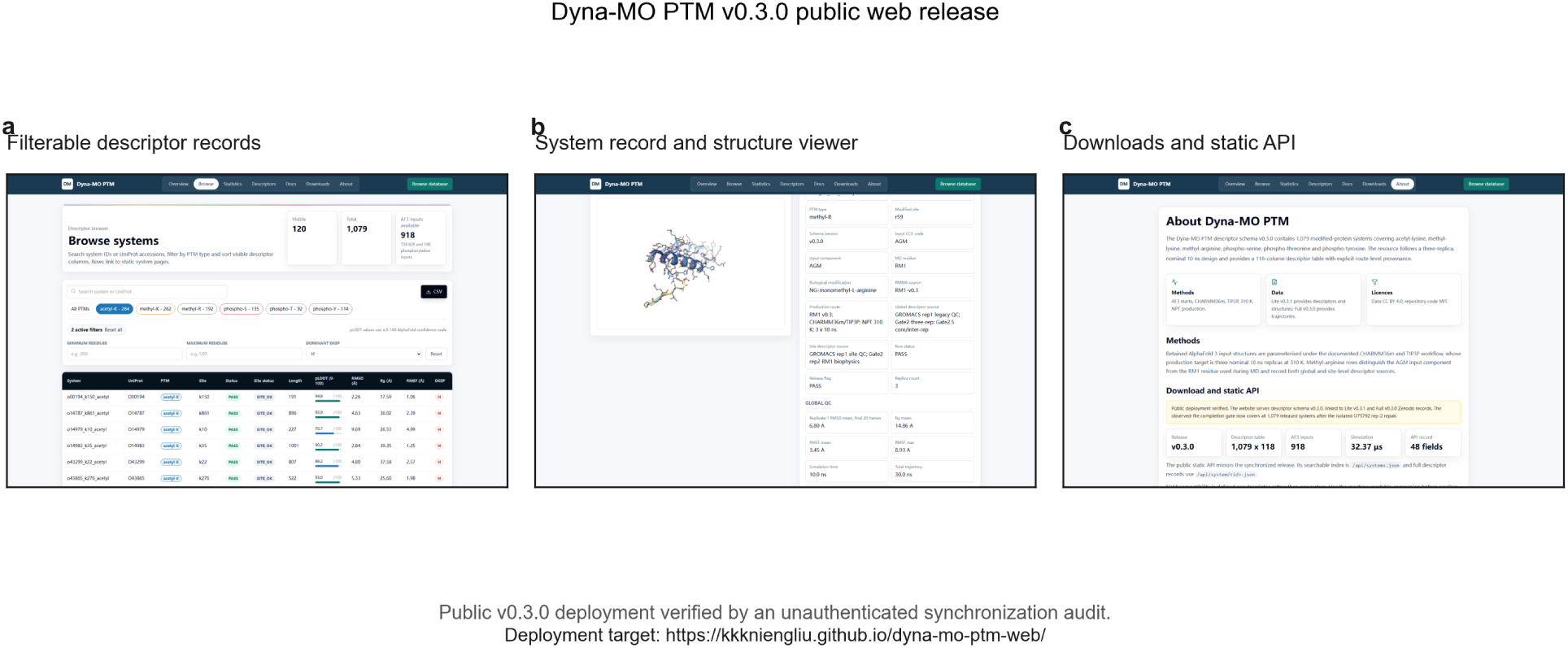
Dyna-MO PTM web interface. The three panels form a discovery, record-inspection and archival-download workflow in the synchronized public v0.3.0 build. (a) The filterable browser identifies records by PTM class and descriptor fields. (b) The per-system page combines provenance and descriptors with an NGL view of the AF3 starting structure. (c) The download and static-API page exposes the master CSV, JSON index and per-system records before users retrieve versioned files from Zenodo. The amber notice states that the observed-file completion gate covers all 1,079 released systems.

### Descriptor-table column layout

The 118 columns of `master_table_all_v2.csv` are summarized here in five user-facing groups. The column dictionary defines eleven functional schema blocks.

- Provenance fields include system identifiers, UniProt accession, PTM type and modified-residue index. Seven correction-provenance fields distinguish input component, MD residue, biological modification, source, production route and descriptor sources for the 192 released methyl-R rows. These new fields are blank for the other classes, whose established provenance fields were preserved during reconciliation.
- AF3 confidence fields have distinct coverage. Starting-PDB B-factor summaries (`plddt_mean`, `plddt_median`, `plddt_min`, `plddt_p10` and `site_plddt`) are populated for all 738 K/R rows. The seven-column raw-output block (`af3_ptm`, `af3_ranking_score`, `af3_fraction_disordered`, `af3_has_clash`, `af3_pae_mean`, `af3_pae_at_site`, `af3_pae_site_local_8A`) and `af3_source` are populated for 918 rows. The context field `local_plddt_8A` is available for 1,029 rows, and `site_plddt_wt` is populated for 161 phosphorylation references.
- Global trajectory fields include Cá-RMSD, Rg, per-residue RMSF and drift slopes. RMSF amplitudes are descriptors rather than global gate criteria. K/R primary columns are the GROMACS rep-1 values used by the route defined above. The explicit K/R three-replica fields separately average full-trajectory RMSD and Rg means and the final-50% equilibrium RMSD mean. Phosphorylation primary columns are three-replica means inherited from DynaMo-phos and mirrored into the explicit `_3rep` aliases in schema v0.3.0.
- Site-biophysics fields include site SASA, the K/R-D/E ionic-contact proxy, χ₁ and χ₂ rotamer states, and the dominant DSSP class. K/R values were computed on rep 2. For phosphorylation systems, three-replica means populate the source-route SASA, contact, rotamer-occupancy and raw DSSP-fraction fields, from which canonical dominant-state aliases were retained. The canonical `site_ss_H_pct`, `site_ss_E_pct`, `site_ss_C_pct` and `site_ss_stability` fields remain blank for these rows.
- Site structural-context fields include terminal distances, burial class and local pLDDT. The K/R subset has the complete ten-column block. Among phosphorylation rows, `is_terminal_lt10`, `burial_class` and `local_plddt_8A` are populated for 291 of 341 systems. `site_plddt_wt` is populated separately for 161 reverted-reference systems, whereas the remaining K/R-specific coordinate-neighbour fields are blank.

Full per-column definitions are in master_table_README.md.

`schema_version` is the version field for downstream parsers; the current table reports `v0.3.0`. This release replaces the methyl-R trajectory descriptors after deterministic conversion from the AGM input component to the RM1 MD residue and excludes 16 corrected systems classified `WARNING`. It preserves the 887 non-methyl-R rows and their order, apart from the declared schema-version transition, and adds seven route-provenance fields. The locked v0.2.8 input table is retained as a checksum-labelled backup.

`cv_fold` provides a deterministic five-fold split at UniProt level. The 1,079 systems map to 1,025 accessions. Of these, 975 occur once and 50 occur two or three times. All rows sharing an accession remain in the same fold. Fold sizes are 209, 211, 218, 221 and 220 systems. This split prevents exact accession leakage but does not control remote homology.

### Dataset composition

Per-PTM counts and CCD codes are given in Table 3.

**Table 3.** Dataset composition, force-field residue codes and modification state as parametrised.

| PTM type | Count | Input component | MD residue | State represented | Release stratum |
| --- | --- | --- | --- | --- | --- |
| Acetyl-lysine | 284 | ALY | ALY | ε-amine acetylated and neutral | K/R subset |
| Monomethyl-lysine | 262 | MLZ | MLZ | monomethylated; dimethylated and trimethylated states not included | K/R subset |
| Monomethyl-arginine | 192 | AGM | RM1 | $N^G$-monomethyl-L-arginine, net charge +1 | K/R subset |
| Phospho-serine | 135 | SEP | SEP | dianionic phosphate at pH 7.4 | DynaMo-phos archive |
| Phospho-threonine | 92 | TPO | TPO | dianionic phosphate at pH 7.4 | DynaMo-phos archive |
| Phospho-tyrosine | 114 | PTR | PTR | dianionic phosphate at pH 7.4 | DynaMo-phos archive |
| Total | 1,079 | | | 32.37 μs completed sampling | Combined release |

The release covers monomethylated lysine and arginine; higher lysine methylation and arginine dimethylation states are outside its scope.

## Data Overview

Figure 1 summarises the 1,079 systems, their protein-length range and the K/R retention funnel. In the common three-replica columns, median final-50% Cα-RMSD is 3.29 Å (IQR 1.94–5.89 Å) and median full-trajectory Rg is 23.01 Å (IQR 19.66–29.19 Å). The route-specific mean Cα-RMSF field has a median of 1.37 Å (IQR 0.88–2.29 Å; Figure 3). These distributions describe the released records rather than matched modification effects.

## Technical Validation

Technical validation tests the internal consistency of the released chemistry, files, quality fields and descriptor transformations. The checks cover release reconciliation, route comparability, starting-structure coverage and sampling limits.

### Release reconciliation and trajectory completion

The K/R release contains 738 systems classified `PASS`. A separate completion gate retained the 341 phosphorylation systems, whose blank K/R flag fields have no `PASS` meaning. The observed-file audit found three 101-frame trajectories spanning 0–10 ns for every released system.

The audit initially found that rep 2 of `o75792_s18_phospho` ended at 3.8 ns. We continued an isolated copy to 10 ns and verified its trajectory, energy, continuity and processing records. The affected per-replica and three-replica descriptors were then regenerated before the release row was updated. A separate hash chain preserves the short trajectory and pre-repair master.

### Uniform per-replica diagnostic

The uniform workflow classified 3,121 of 3,237 replicas as PASS, 102 as WARNING and 14 as FAIL under the inherited K/R engineering thresholds. At system level, all three replicas passed for 988 of 1,079 systems; the worst replica was WARNING for 78 systems and FAIL for 13.

Within the released K/R subset, rep 1 yielded 737 PASS and one marginal WARNING, rep 2 yielded 699 PASS, 37 WARNING and two FAIL, and rep 3 yielded 691 PASS, 42 WARNING and five FAIL. Thus, 670 of 738 K/R systems had three PASS replicas, while 62 had a worst label of WARNING and six had a worst label of FAIL. The single rep-1 reclassification, `f0nhv6_k25_methyl`, crossed only the relative-fluctuation warning boundary when recalculated from the deposited trajectory (0.303, compared with 0.297 in the source gate series). The exact source-route release status remains PASS. Of direct relevance to site-biophysics reuse, 39 K/R rep-2 trajectories received a non-PASS diagnostic, comprising 37 WARNING and two FAIL.

For phosphorylation, 994 of 1,023 replicas passed the same diagnostic, 22 were WARNING and seven were FAIL. All three passed in 318 of 341 systems; the worst replica was WARNING in 16 and FAIL in seven. These labels identify trajectories for sensitivity analysis. A threshold crossing can reflect large domain motion as well as undesirable drift, so it is not by itself evidence of unfolding or file corruption. Supplementary Table S2 and the companion CSVs provide every replica value and issue code.

### Cross-route descriptor comparability

The K/R and phosphorylation records form two provenance strata. Primary K/R global fields come from rep 1, whereas populated phosphorylation values are inherited three-replica summaries. K/R site-biophysics fields come from rep 2; their phosphorylation counterparts again use inherited three-replica summaries. The column dictionary marks each of these fields as route-specific.

All 1,079 systems have three-replica RMSD and Rg fields. Among the 738 K/R systems, the median absolute difference between primary and three-replica RMSD is 0.323 Å (Spearman ρ = 0.969). For Rg, the corresponding difference is 0.094 Å (ρ = 0.999). These values support use of the common RMSD and Rg summaries. Comparisons involving RMSF or site-biophysics descriptors should retain the recorded production and analysis routes.

### Sensitivity of the K/R release gate

The threshold audit began with all 1,089 K/R systems that completed the original production workflow. Full-precision rep-1 GROMACS series were recovered for the 787 acetyl-K and methyl-K systems. For methyl-R, the 208 corrected RM1 series replaced their superseded AGM-route values. The remaining 94 completed methyl-R systems lacked corrected chemistry and were held out at every threshold scale, leaving 995 chemistry-eligible systems for numerical perturbation. Reapplying the original cutoffs reproduced 738 PASS systems exactly. Tightening all absolute cutoffs by 20% retained 670, whereas relaxing them by 20% retained 799. The corresponding acetyl-K, methyl-K and methyl-R counts were 255, 246 and 169 under the tighter gate; 284, 262 and 192 under the original gate; and 309, 292 and 198 under the relaxed gate. This analysis shows how strongly release size depends on the inherited engineering cutoffs without treating any scale as a biological boundary.

### Solvent padding and periodic images

Preparation logs identify 731 K/R systems built with 0.90 nm padding, five with 0.75 nm and two with 0.60 nm. We evaluated the minimum macromolecule-to-periodic-image distance at every stored frame of all 21 replicas belonging to the seven systems with reduced padding. Nineteen replicas remained at or above the 1.2 nm non-bonded cutoff throughout. Two replicas each contained one closer saved frame: `q13620_k698_acetyl` rep 2 reached 1.182 nm at 5.7 ns and `q8nb14_r535_methyl` rep 3 reached 1.190 nm at 1.6 ns. No 0.60 nm-padding replica crossed the cutoff. Because coordinates were saved every 100 ps, the duration of either approach cannot be resolved more finely. Both systems are retained with an explicit periodic-image warning in the audit table. The first warning applies to the replica used for site-biophysics descriptors; the second applies to a replica contributing to three-replica summaries.

### Truncation-created termini

We combined the locked K/R core coordinates with UniProt sequence lengths retrieved on 21 August 2026. At least one retained boundary was truncation-created in 561 of 738 systems. Twenty-nine modified sites were fewer than 10 residues from an artificial terminus, and 16 were fewer than five residues away. The system-level audit distinguishes natural from truncation-created N and C termini and records the retrieval URLs. Because the original UniProt sequence snapshot was not retained, this reconstruction uses current canonical or specified-isoform lengths rather than claiming an historical sequence snapshot.

### RM1 topology and production checks

For every RM1 topology, the audit found one target residue, the required atoms, a +1 charge and resolvable bonded terms. All 624 corrected replicas also passed file, log, trajectory and post-processing checks before reconciliation of the 192-system release subset. This establishes internal consistency for the recorded CHARMM36m route, but not independent QM, torsional or experimental validation of RM1.

### Starting-structure and schema checks

Raw AF3 confidence outputs matched 918 of the 1,079 release rows through exact identifiers or verified truncation offsets. The other 161 rows, all from phosphorylation batch 2, contain null values because their raw outputs were not retained. Site-window pLDDT should be interpreted conditional on the K/R input filter.

All 1,079 rows have canonical dominant χ₁ and DSSP labels. Field-availability masks, rotamer vocabularies and units were checked against the frozen schema. Canonical SASA is in nm², `site_ss_stability` is on a 0–100 scale, and the ionic-contact quantity reports a protein-wide proxy.

### Association between AF3 confidence and finite-window relaxation

We assessed whether confidence in the selected AF3 input was associated with four finite-window descriptors in the 738 K/R systems, the subset with a common input filter and complete starting-PDB pLDDT summaries. Mean pLDDT was inversely associated with final-20% backbone RMSD (Spearman ρ = −0.634). The association remained within each PTM class (ρ = −0.604 for acetyl-K, −0.637 for methyl-K and −0.621 for methyl-R) and after rank-residual adjustment for protein length and PTM class (*r* = −0.577). Site pLDDT showed a weaker association with the same RMSD descriptor (ρ = −0.428; adjusted *r* = −0.345). The complete eight-comparison table, covering RMSD level and slope, Rg and the high-flexibility fraction, is provided in `af3_descriptor_associations_v0.3.0.csv` with pairwise-complete sample counts and class-specific estimates. These descriptive relationships are conditional on the selected and, where necessary, truncated AF3 inputs. They do not establish causal error propagation or calibrate uncertainty in the MD descriptors.

### Within-resource paired pipeline diagnostic

Across the six reverted-to-canonical pairs, mean phospho-minus-canonical differences were +0.07 Å for backbone RMSD, +0.02 Å for Rg and +0.03 Å for site Cα RMSF. Wilcoxon signed-rank p values exceeded 0.4 (Supplementary Figure S1). The six pairs establish analyzable matched trajectories and serve as a pipeline diagnostic rather than an effect or equivalence estimate.

### Early-versus-late block diagnostic

All 284 acetyl-K rep-1 trajectories extended to 100 ns. The non-overlapping 0–10 ns and 90–100 ns blocks each contained 101 frames. Between these blocks, backbone RMSD increased by 2.70 Å on average (RMSE 3.98 Å; Pearson *r* = 0.860), whereas Rg decreased by 0.53 Å (RMSE 1.72 Å; *r* = 0.979; Supplementary Figure S2). The persistent differences provide no evidence of convergence or equivalence in this one-replica acet-K diagnostic.

## Usage Notes

Stable system identifiers link the descriptor table, starting structures, trajectories and provenance manifests.

### Record-level user journey

A user can begin with the browser or JSON index to filter systems by PTM type and descriptor. The selected record then exposes its production route, quality-control fields and starting-structure availability. After inspecting the record and structure, the user can download the lite record for tables and manifests, retrieve the corresponding trajectory files from the full record, and recompute descriptors while retaining the published identifiers and checksums.

### Descriptor access and aggregation

Complete 118-column records are available in `master_table_all_v2.csv`. The smaller `/api/ systems.json` file supports search, and `/api/system/<system_id>.json` provides individual records. The field-level companion and `/api/descriptor-compatibility.json` expose the same route-aware reuse contract without adding a row-level label to the frozen master schema. K/R `PASS` labels come from the rep-1 gate; phosphorylation rows were retained by completion-based curation. Because their aggregation histories differ, pooled analyses should use the three-replica RMSD and Rg aliases where possible and retain source-route strata. Missing AF3 and site-context values should remain missing rather than being replaced with zero.

The uniform-QC companion supports stricter reuse filters without changing the master table. Users can select the 988 systems with three diagnostic PASS replicas or choose route- and replica-specific criteria. For K/R site SASA, contacts, rotamers and DSSP, `rep2_diagnostic_status` identifies the quality label of the analysed replica; analyses sensitive to global drift should exclude or separately examine its 39 non-PASS records.

### Descriptor families and interpretation

The descriptor groups address different reuse questions and should not be treated as interchangeable measures of stability or PTM effect (Table 5). Cross-route screening should begin with the common three-replica RMSD and Rg fields. Site-biophysics and structural-context descriptors can then address local questions within analyses that retain the relevant production-route strata.

**Table 4.** Functional comparison with representative PTM-aware and general MD resources.

| Resource | Primary scope | MD product or output | Structured comparison layer and principal reuse |
| --- | --- | --- | --- |
| Dyna-MO PTM | Six protein PTM classes | Three deposited 10 ns replicas for each of 1,079 systems | A 118-column descriptor and provenance table for filtering, reanalysis and method evaluation |
| ATLAS<sup>11</sup> | General protein flexibility | Standardised protein simulations downloadable at several processing levels | Per-protein dynamics summaries, search and programmatic access |
| mdCATH<sup>12</sup> | Protein domains across temperatures | Coordinates and forces from replicated all-atom simulations | Derived metadata and loaders for biophysical analysis and data-driven modelling |
| GPCRmd<sup>13</sup> | G-protein-coupled receptors | Curated and community-shared GPCR simulations | Interactive visualisation and comparative analysis of receptor dynamics |
| MDRepo<sup>14</sup> | Community-contributed biomolecular MD | Deposited simulations from multiple contributors | Search, large-scale retrieval, visualisation and cloud-backed analysis |
| PTMdyna<sup>15</sup> | User-specified PTM comparisons | Server-generated unmodified and modified conformational ensembles | Case-specific comparison of structural and dynamical properties |
| Vienna-PTM<sup>16</sup> | PTM system preparation | Simulation-ready structures, topologies and force-field parameters | Preparation and parametrisation rather than a cross-system trajectory archive |

**Table 5.** Recommended interpretation of the principal descriptor families.

| Descriptor family | Supported interpretation | Boundary of interpretation |
| --- | --- | --- |
| Global RMSD, Rg and RMSF | Finite-window structural relaxation, molecular extent and fluctuation screening | Not evidence of thermodynamic stability, convergence or kinetics |
| Site SASA and burial context | Accessibility and local structural context of the recorded site | Not a modification probability, enzyme-recognition score or reaction rate |
| Rotamer occupancy and DSSP | Local conformations sampled during the deposited window | Not an equilibrium population or transition model |
| K/R–D/E ionic-contact proxy | Frequency of recorded geometric near contacts | Not a salt-bridge energy, electrostatic free energy or binding free energy |
| AF3 confidence fields | Confidence and selection context for the starting model | Not proof of MD accuracy or a calibrated uncertainty on trajectory descriptors |

### Truncation boundaries and solution conditions

Distances to the retained input termini are reported in `dist_to_nterm` and `dist_to_cterm`, and `is_terminal_lt10` identifies sites fewer than 10 residues from either boundary. This broad flag applies to 60 of the 738 K/R systems; 32 are fewer than five residues from a retained boundary. The separate artificial-terminus audit narrows these counts to 29 and 16, respectively. Truncation-created boundaries were simulated as standard charged termini rather than capped peptide bonds, so users studying local electrostatics or solvent exposure near a flagged site should filter or model those systems separately. The solvent contains only the Na^+^ or Cl^−^ ions required for charge neutrality. It does not represent a prescribed physiological bulk-salt concentration.

The periodic-image audit flags `q13620_k698_acetyl` and `q8nb14_r535_methyl` because one stored frame in one replica of each system was 0.018 nm or 0.010 nm inside the 1.2 nm cutoff. Analyses sensitive to occasional surface-image proximity should exclude these systems or repeat them in a larger solvent box.

### Appropriate and inappropriate reuse

Suitable uses include descriptor reproduction, route-stratified ranking, method development and selection of systems for longer simulations. The reported occupancies, dominant states and exposure measures describe the sampled 10 ns windows. Estimating folding rates, transition kinetics, equilibrium free energies or population-level substrate preferences requires longer, purpose-designed simulations. Quantifying modification effects additionally requires matched unmodified trajectories. RM1 applications outside the recorded protocol should include a target-specific force-field assessment.

Three workflows provide concrete starting points for reuse.

1. For candidate selection, filter by PTM class, burial, solvent exposure or local structure, apply the relevant route and replica quality-control fields, inspect the retained coordinates, and nominate systems for longer simulations.
2. For descriptor baselines, predefine compatible fields from Table 2, retain production-route strata, report missingness, and use accession-level or task-specific sequence-cluster splits before fitting or comparing models.
3. For external validation of trajectory-analysis or generative methods, select held-out systems, compare model outputs with deposited coordinates using prespecified observables, and report validity failures separately from numerical agreement. The 10 ns references support finite-window structural comparisons, not kinetic validation.

### Sequence splits and missing values

Sequence-based benchmarks should supplement the deterministic UniProt-level folds with task-specific cluster splits to control remote homology. Site-context descriptors are available for 291 of 341 phosphorylation systems; the remaining 50 are listed in `phos_site_context_wt.missing.md`.

The CHARMM36m and TIP3P combination defines the force-field and water-model scope of the trajectories^31,32^.

### Additional limitations

Cross-route pooled analyses should retain the provenance strata described in Technical Validation. `af3_source` separates 813 exact matches, 105 verified numbering remaps and 161 batch-2 rows without retained raw AF3 outputs. The source records preserve the final site identities but not the database snapshots, evidence filters or conflict rules used during initial assembly. The archived K/R AF3 runner identifies its v3.0.0 code base and database filenames but does not retain a model-parameter checksum or a dated PDB mmCIF snapshot. The release should therefore be treated as a curated PTM resource rather than a proteome-representative cohort.

## Data Availability

The Dyna-MO PTM v0.3.1 lite record contains the v0.3.0 descriptor table, column dictionary, retained starting structures, manifests, supplementary tables, field-level compatibility metadata and analysis scripts (Zenodo, https://doi.org/10.5281/zenodo.22059934)^30^. The companion full record contains the 133.70 GB trajectory payload, a 3,237-row replica manifest, topology companions, reference coordinates and hash-chain audit evidence (Zenodo, https://doi.org/10.5281/zenodo.22049767)^29^. Both records are publicly accessible under CC BY 4.0. The lite record does not redistribute CHARMM36m files. It records the official July 2022 source and checksum and provides the project-authored RM1 installation and validation scripts; users obtain the base force field from the MacKerell Lab. The synchronized v0.3.0 discovery browser is available at https://kkkniengliu.github.io/dyna-mo-ptm-web/.

## Code Availability

Data preparation, MD orchestration, quality control, descriptor extraction and figure generation are scripted in the source repository under the MIT license^28^. The v0.3.1 lite Zenodo record archives the analysis and release code used for the v0.3.0 dataset, including the AGM-to-RM1 conversion, RM1 installation, topology-validation and P1 reuse-metadata scripts. Scripts require Python 3.10 or later. The analysis code uses MDTraj, NumPy, pandas, SciPy and Matplotlib^33–37^. Release statistics were generated with Python 3.13.5, NumPy 2.3.3, pandas 2.2.3 and SciPy 1.16.2. The corrected RM1 and GROMACS quality-control bridge workflows used Python 3.11.15, MDTraj 1.11.1, NumPy 2.3.5 and SciPy 1.16.3. Exact package versions for the legacy workflows and historical Matplotlib version were not retained. GROMACS versions and supporting runtime evidence are recorded by production route in the Methods and archived run manifests.

## Author Contributions

K.L. (Kaining Liu) contributed to conceptualization, methodology, software, validation, formal analysis, investigation, data curation, original-draft writing and visualization. Q.Q. (Qiuting Qian) contributed resources from the DynaMo-phos phosphorylation archive, methodology, investigation, data curation, review and editing. J.P. (Jiahua Peng), D.M. (Dongge Ma), Y.Y. (Yifei Yao) and J.Z. (Jingjing Zhao) contributed to investigation, data curation, review and editing. Y.C. (Ying Chi) contributed to conceptualization, supervision, project administration, funding acquisition, review and editing.

## Acknowledgements

We thank the high-performance computing facilities of the Zhejiang University–University of Edinburgh Institute and the Second Affiliated Hospital of Zhejiang University School of Medicine for computational resources.

## Ethics statement

This computational study involved no human participants, animals or identifiable personal data.

