## Supplementary Information for "Molecular dynamics descriptors for 1,079 post-translationally modified protein systems"

#### Contents


- Table S1. Cross-route descriptor comparability contract.
- Table S2. Uniform per-replica trajectory diagnostic.
- Table S3. Sensitivity of the K/R release thresholds.
- Table S4. Periodic-image audit for reduced-padding systems.
- Table S5. Truncation-created termini in the K/R subset.
- Figure S1. Within-resource paired pipeline diagnostic for six phospho-S/T/Y systems.
- Figure S2. Acetyl-K early-versus-late block diagnostic for 284 systems, rep 1.


#### Table S1

Table S1. Cross-route descriptor comparability contract.

| Descriptor family | K/R source | Phosphorylation source | Completeness | Recommended reuse |
|---|---|---|---|---|
| `rmsd_mean_3rep_A` | Mean of three full-trajectory replica means | Alias of inherited three-replica mean | 1,079/1,079 | Preferred common RMSD summary, with source route retained |
| `rmsd_equil_mean_3rep_A` | Mean of three final-50% replica means | Alias of inherited three-replica equilibrium mean | 1,079/1,079 | Preferred common equilibrium-segment RMSD summary |
| `rg_mean_3rep_A` | Mean of three full-trajectory replica means | Alias of inherited three-replica mean | 1,079/1,079 | Preferred common Rg summary |
| Primary RMSD and Rg fields | GROMACS rep 1; release-route windows | Inherited three-replica summaries | 1,079/1,079 | Route-specific QC and descriptive use only |
| Primary RMSF fields | K/R rep 1; additional K/R-only three-replica variants | Inherited three-replica summary | Route-dependent | Do not pool across routes as one measurement definition |
| Site RMSD and site RMSF | K/R rep 1 | Unavailable | 738/1,079 | K/R-only diagnostic |

| | | | | |
|---|---|---|---|---|
| Site SASA and ionic-contact proxy | K/R rep 2; GROMACS or MDTraj by subtype | Inherited three-replica means | 1,079/1,079 | Route-specific and descriptive |
| Rotamer and dominant DSSP labels | K/R rep 2 | Inherited three-replica summaries | 1,079/1,079 | Category filtering; do not infer PTM effects from pooled classes |
| Canonical DSSP percentages and stability | K/R rep 2 | Unavailable | 738/1,079 | K/R-only occupancy summaries |
| AF3 raw-output confidence fields | Retained raw output | Retained batch-3 raw output | 918/1,079 | Analyse with `af3_source`; do not impute missing batch-2 values |

The three `_3rep_A` fields align replica aggregation across the two source routes. Software, fitting procedures, analysis windows and release gates nevertheless remain route-specific. Exact definitions and field availability are given in the master-table dictionary.

#### Table S2

Table S2. Uniform per-replica trajectory diagnostic.

| Route and unit | Replica | Number assessed | PASS | WARNING | FAIL |
|---|---|---|---|---|---|
| K/R replicas | 1 | 738 | 737 | 1 | 0 |
| K/R replicas | 2 | 738 | 699 | 37 | 2 |
| K/R replicas | 3 | 738 | 691 | 42 | 5 |
| K/R systems, worst of three replicas | N/A | 738 | 670 | 62 | 6 |
| Phosphorylation replicas | 1 | 341 | 331 | 9 | 1 |
| Phosphorylation replicas | 2 | 341 | 326 | 11 | 4 |
| Phosphorylation replicas | 3 | 341 | 337 | 2 | 2 |
| Phosphorylation systems, worst of three replicas | N/A | 341 | 318 | 16 | 7 |
| All replicas | N/A | 3,237 | 3,121 | 102 | 14 |
| All systems, worst of three replicas | N/A | 1,079 | 988 | 78 | 13 |

The same GROMACS selections and the exact legacy K/R engineering thresholds were applied diagnostically to every released replica. These labels supplement the source-route release decisions; they do not redefine them. The companion CSV provides all 3,237 per-replica records and a 1,079-row system summary. A system-level status is the worst label among its three replicas. Among the K/R records, 39 rep-2 trajectories used for site-biophysics descriptors were non-PASS, comprising 37

WARNING and two FAIL.

#### Table S3

Table S3. Sensitivity of the K/R release thresholds.

| Threshold scale | Acetyl-K PASS | Methyl-K PASS | Methyl-R PASS | Total PASS | WARNING | FAIL |
|---|---|---|---|---|---|---|
| 0.8× | 255 | 246 | 169 | 670 | 182 | 143 |
| 1.0× | 284 | 262 | 192 | 738 | 176 | 81 |
| 1.2× | 309 | 292 | 198 | 799 | 159 | 37 |

The audit covers 995 chemistry-eligible completed systems. A further 94 completed methyl-R systems lacked corrected RM1 chemistry and remained ineligible at every scale. The relative RMSD fluctuation threshold of 0.30 was held fixed; all absolute RMSD, slope and Rg-CV warning and failure cutoffs were multiplied by the scale shown.

#### Table S4

Table S4. Periodic-image audit for the seven K/R systems prepared with less than 0.90 nm solvent padding.

| System | PTM type | Padding (nm) | Minimum distance over three replicas (nm) | Stored frames below 1.2 nm | Affected replicas |
|---|---|---|---|---|---|
| `o14920_r265_methyl` | Methyl-R | 0.75 | 4.052 | 0 | 0 |
| `p51784_k484_acetyl` | Acetyl-K | 0.75 | 1.372 | 0 | 0 |
| `p51784_r630_methyl` | Methyl-R | 0.60 | 1.266 | 0 | 0 |
| `q13620_k698_acetyl` | Acetyl-K | 0.75 | 1.182 | 1 | 1 |
| `q8iy17_k684_acetyl` | Acetyl-K | 0.60 | 1.451 | 0 | 0 |
| `q8nb14_r535_methyl` | Methyl-R | 0.75 | 1.190 | 1 | 1 |
| `q9ujz1_k323_methyl` | Methyl-K | 0.75 | 1.341 | 0 | 0 |

All 101 saved frames were assessed in each of the 21 replicas. `q13620_k698_acetyl` rep 2 reached 1.182 nm at 5.7 ns, and `q8nb14_r535_methyl` rep 3 reached 1.190 nm at 1.6 ns. The 100 ps output interval does not resolve how long either approach persisted.

#### Table S5

Table S5. Truncation-created termini in the 738-system K/R release.

| PTM type | Systems | At least one artificial terminus | Site <10 residues from artificial terminus | Site <5 residues from artificial terminus |
|---|---|---|---|---|
| Acetyl-K | 284 | 225 | 12 | 6 |
| Methyl-K | 262 | 168 | 10 | 8 |
| Methyl-R | 192 | 168 | 7 | 2 |
| Total | 738 | 561 | 29 | 16 |

Artificial boundaries were reconstructed by comparing the locked structural-core coordinates with UniProt canonical or specified-isoform lengths retrieved on 21 August 2026. The original sequence snapshot was not retained, so the companion table records the retrieval source and should be interpreted as a current-sequence reconstruction.

#### Figure S1

Within-resource paired pipeline diagnostic

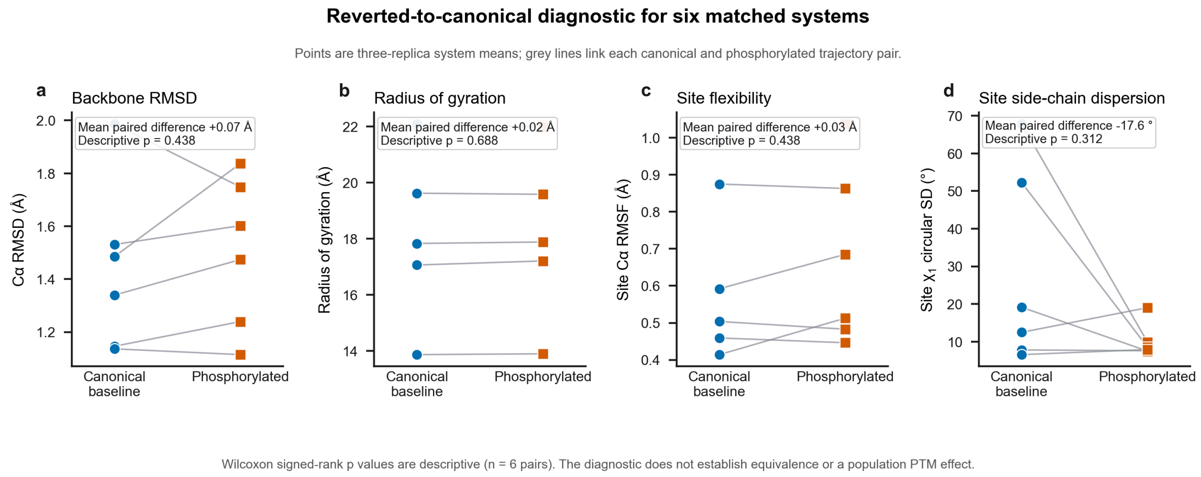


Figure S1. Paired pipeline diagnostic for six phospho-S/T/Y systems. Each AF3-seeded phosphorylated structure was paired with a 3 × 10 ns trajectory in which SEP, TPO or PTR was reverted to Ser, Thr or Tyr. Both members of a pair followed the same CHARMM36m and TIP3P workflow. Panels a–d show backbone Cα-RMSD relative to frame 0, Rg, site Cα-RMSF and modified-site $\chi_1$ circular standard deviation. Lines join the three-replica means within each pair. Wilcoxon signed-rank p values are descriptive. With only six convenience pairs, non-significant results cannot establish equivalence or exclude a phosphorylation effect.

### Figure S2

Acetyl-K early-versus-late block diagnostic

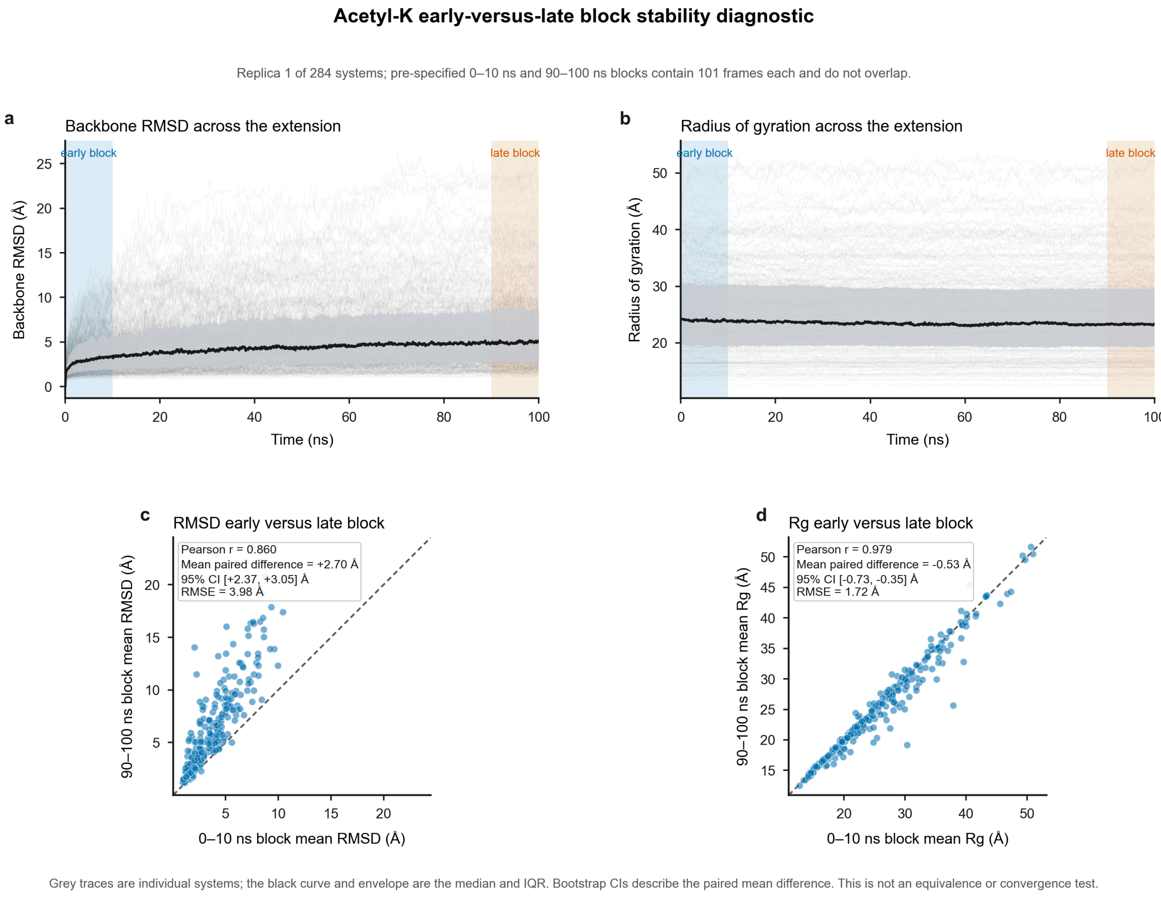


Figure S2. Early-versus-late diagnostic for rep 1 of 284 acetyl-K systems. Panels a and b show backbone RMSD and Rg over 100 ns. Grey traces represent individual systems; the black line and shaded region show the median and interquartile range. Blue and orange mark the non-overlapping 0–10 ns and 90–100 ns blocks. Panels c and d compare the block means. For RMSD, $r = 0.860$, the paired difference is +2.70 Å (95% CI +2.37 to +3.05 Å) and RMSE is 3.98 Å. For Rg, $r = 0.979$, the paired difference is −0.53 Å (95% CI −0.73 to −0.35 Å) and RMSE is 1.72 Å. The comparison was not designed as a convergence or equivalence test.
